# Age-related intrinsic functional connectivity changes of locus coeruleus from childhood to older adults

**DOI:** 10.1101/2021.10.10.463574

**Authors:** Inuk Song, Joshua Neal, Tae-Ho Lee

**Affiliations:** Department of Psychology, Virginia Tech; School of Neuroscience, Virginia Tech

**Keywords:** locus coeruleus, distractibility, neurodevelopment, functional connectivity

## Abstract

The locus coeruleus is critical for selective information processing by modulating brain’s connectivity configuration. Increasingly studies have suggested that LC controls sensory inputs at the sensory gating stage. Furthermore, accumulating evidence has examined that young children and older adults are more prone to distraction and filter out irrelevant information less efficiently, possibly due to the impaired LC connectivity. However, the LC connectivity pattern across the life span is not fully examined yet, hampering our ability to understand the relationship between LC development and the distractibility. In this study, we examined the intrinsic network connectivity of the LC using resting-state fMRI from the enhanced NKI dataset with wide-range age samples. Based on LC-seed functional connectivity maps, we examined the age-related variation in the LC connectivity with a quadratic model. The analyses revealed two connectivity patterns explicitly. The sensory-related brain regions showed a positive quadratic age effect (u-shape), and the frontal regions for the cognitive control showed a negative quadratic age effect (inverted u-shape). Our results imply that such age-related distractibility is possibly due to the impaired sensory gating by the LC and the insufficient top-down controls by the frontal regions. We discuss the underlying neural mechanisms and limitations of our study.

## INTRODUCTION

The locus coeruleus (LC) is a small nucleus located deep in the brainstem and a major source of norepinephrine. The LC releases norepinephrine to almost the entire brain throughout its efferent projections, thereby the LC is one of the primary brain regions critical for selective information processing by changing the brain’s configurations at both phasic and tonic levels (Aston-Jones and Cohen, 2005; Berridge and Waterhouse, 2003; Carter *et al*., 2010; Chen, 2002; Clewett *et al*., 2018; Coronel-Oliveros *et al*., 2021; Lee *et al*., 2018; Lee *et al*., 2020b; Lee *et al*., 2014; Mather and Harley, 2016). Recent studies consistently have suggested that the LC functionally controls sensory inputs at the early sensory gating stage by changing the brain’s connectivity configurations (Devilbiss *et al*., 2012; McBurney-Lin *et al*., 2019; Waterhouse and Navarra, 2019). For instance, the stimulated LC changes its neural communication with the basolateral nucleus of the amygdala (Fast and McGann, 2017), and thalamus (Devilbiss and Waterhouse, 2011; Rodenkirch *et al*., 2019) that receive the majority of sensory information for the further in-depth cognitive process. Similarly, direct chemo-genetic stimulation of LC immediately changes neural connectivity configurations, especially for the intrinsic networks that mediate the bottom-up sensory process, including the primary sensory and salience networks (Zerbi *et al*., 2019). That is, the LC plays a role in controlling sensory flows in the brain, suggesting that impaired processing selectivity is possibly due to the failure of communication between the LC and sensory regions that introduces the sensory overflows in the brain.

Human imaging studies also imply that the LC system changes the bottom-up process at the early sensory-perceptual stage to prioritize important information. For example, the induced phasic LC activity by arousing or stressful stimuli at the early sensory-perceptual level increases the initial selective attention processes (Lee *et al*., 2018; Lee *et al*., 2020c; Lee *et al*., 2014) and attentional control (Chiew and Braver, 2013; van der Wel and van Steenbergen, 2018). Finally, such initial LC-induced selectivity at the early sensory stage carries over to the late cognitive processing such as memory encoding (Yebra *et al*., 2019), memory consolidation (Clewett *et al*., 2018; Gallant *et al*., 2021), and decision making (Bland, 2012). Thus, the interrupted LC activity has been often examined in individuals with conditions associated with hyperarousal and attentional vigilance such as attention-deficit hyperactivity disorder (ADHD; Bruno *et al*., 2007; Darcq and Kieffer, 2015; Rowe *et al*., 2005), and posttraumatic stress disorder (PTSD; McCall *et al*., 2017; Naegeli *et al*., 2018; Serova *et al*., 2019).

Recent studies also showed that older adults, who are more prone to distraction, exhibit interrupted LC connectivity, (Bachman *et al*., 2021; Clewett *et al*., 2016; Kempadoo *et al*., 2016; Leslie *et al*., 1985; Liu *et al*., 2020; Marcyniuk *et al*., 1986; Mueller *et al*., 2017; Olpe and Steinmann, 1982). For instance, older adults showed the hyper-connectivity of the LC with the primary sensory networks compared to younger adults as well as the hypo-functional coupling with the salience network (Lee *et al*., 2020c), suggesting that the impaired connectivity between the sensory regions and LC induces sensory overflows in the brain and thus the salience network fails to guide attention appropriately. As a result, there is an unnecessary depletion of limited neural resources in the brain leading the executive frontal systems to not maintain goal-directed processes due to irrelevant stimuli that should be ignored earlier.

However, children’s developmental trajectories of LC connectivity have not been characterized yet. Healthy young children also show behavioral propensity to react to irrelevant information combined with heightened impulsivity (Hoyer *et al*., 2021; Kannass *et al*., 2006; Palfrey *et al*., 1985). This distractibility in children is possibly due to their less-developed LC function during early childhood, such that structural studies have demonstrated that structural integrity of the LC increases with age gradually and then declines after the peak (i.e., inverted U-shaped curvilinear trend; Bondareff *et al*., 1981; Bondareff *et al*., 1982; Jacobs *et al*., 2018; Liu *et al*., 2019). Furthermore, studies suggest that brain development occurs first in the primary sensory bottom-up regions from early childhood with a progressively maturing top-down frontal system (Casey *et al*., 2019; Toga *et al*., 2006). Thus, the unbalanced brain development in early childhood between not-fully-developed LC and matured sensory network regions possibly leads children to fail at controlling sensory overflows in the brain.

Considering accumulating evidence indicating that the LC plays an essential role in the brain’s processing selectivity, the overarching objectives of the current study are to provide a full description of LC connectivity pattern across the lifespan from early childhood to older adulthood. Given the brain development findings (Bondareff *et al*., 1981; Bondareff *et al*., 1982; Jacobs *et al*., 2018; Liu *et al*., 2019) and heightened distractibility in early childhood and older adults (Hoyer *et al*., 2021; Kannass *et al*., 2006; Lee *et al*., 2018; Lee *et al*., 2020c; Palfrey *et al*., 1985), we especially hypothesized that children and older adults, compared to younger adults, show increased functional connectivity of the LC with the primary sensory regions (i.e., quadratic or u-shape curve), indicating that those two age groups have unnecessarily higher sensory sensitivity intrinsically even without task-induced activity (i.e., resting-state fMRI). We hypothesized the quadratic age effects because the developmental of structural integrity in the LC follows curvilinear trend (Liu *et al*., 2019) and behavioral performances also follow the inverted U-shape in general (Nomi *et al*., 2017). To this end, we used cross-sectional samples (age-ranged between 8 and 83 years) and examined intrinsic functional connectivity of the LC associated with age changes based on the resting-state fMRI data. We especially examined the intrinsic network connectivity of the LC based on the resting-state fMRI signal as it reflects general intrinsic neural architecture of brain development at the time of the brain scan, rather than a moment-by-moment task-specific neural response (Elliott *et al*., 2019; Lee *et al*., 2017a; b; Lee and Telzer, 2016; Lee *et al*., 2020c).

## MATERIALS AND METHODS

### Data characteristics

The present study was carried out using resting-state fMRI data from the enhanced Nathan Kline Institute (NKI)-Rockland project (Nooner *et al*., 2012). The dataset was initially downloaded through the Mind Research Network’s collaborative informatics and neuroimaging suite (COINS; Landis *et al*., 2016). We only included individuals with full-coverage of both T1 and EPI, and without severe motions (framewise displacement, *FD* > 0.5 mm), resulting in 595 samples (*M* = 39.47 years, *SD* = 20.51, range = 8–83, 63.36% females) samples (see Figure S1 in *Supplementary*). All individual data has been collected in the same scanning protocol with a 32-channel head-coil for the high-resolution structural image (T1-MPRAGE; TR = 1950 ms; TE = 2.52 ms; FA = 9°; 1-mm isotropic voxel; FOV = 256 mm) and EPI image (364 volumes; 2-mm isotropic voxel, 64 slices; TR = 1400 ms; TE = 30 ms; FA = 65°; matrix size = 112 × 112; FOV = 224 mm).

### Preprocessing

Preprocessing was performed using the FSL combined with ICA-AROMA (Pruim *et al*., 2015) and ANTs (Avants *et al*., 2014), including skull stripping and tissue mask segmentation (CSF/WM/GM) after bias-field correction for structural images, and first ten-volumes cut, motion correction, slice-timing correction, intensity normalization, regressing out CSF/WM with individually segmented masks, ICA-denoising (corrected mean FD = 0.02 mm, range = 0.01 – 0.14 mm; Figure 1B) and registration to standard MNI 2-mm brain template for functional images. To avoid possible signal mixture of LC region with neighboring regions such as periaqueductal gray or ventral tegmental area, we skipped signal smoothing step.

**Figure 1.**
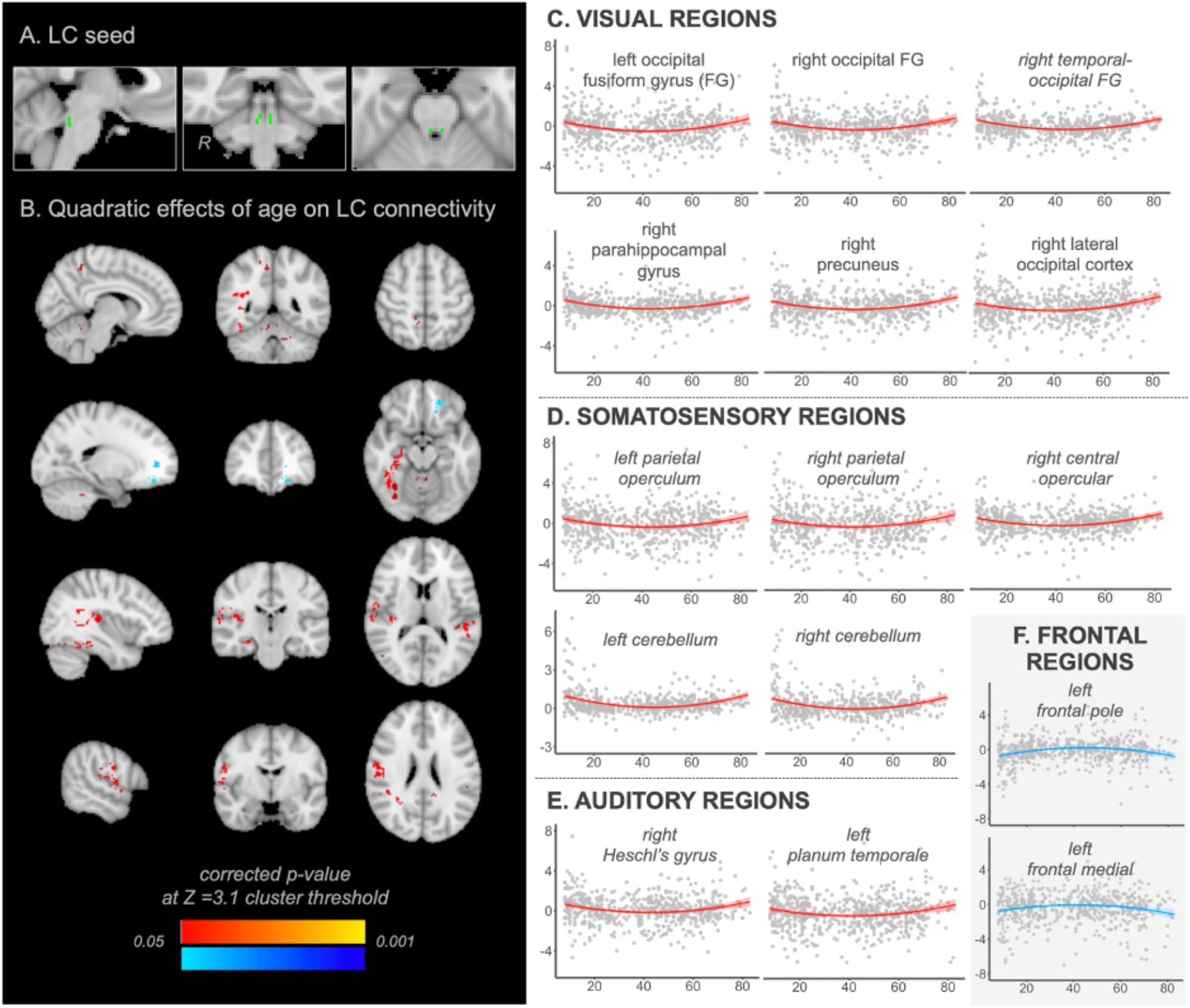
(A) LC seed mask (B) whole brain maps showing a quadratic effect of age on LC connectivity. Regions in (C) the visual network (D) somatosensory network (E) auditory network and (F) frontal network. Y-axis: LC connectivity strength with each region; X-axis: age.

### Whole-brain multiple regression analysis for age-related changes in LC connectivity

We first extracted the mean time-series of LC activity from the preprocessed image on each individual’s non-smoothed native space using a standard structural LC mask (Figure 1A; Keren *et al*., 2009). Using this LC time course, a multiple regression analysis was then performed to estimate individual level LC-seed functional connectivity maps. Finally, changes of LC connectivity with age were estimated at the whole-brain level using a multiple regression model. Consistent with our main hypothesis, we examined the age-related variation in the LC connectivity with a quadratic model (i.e., *age2* and *age):*

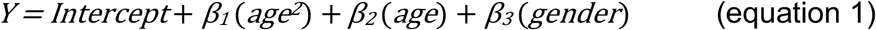

In the model, we included *gender* in the design matrix as nuisance regressors to attenuate gender effects. The group-level whole-brain connectivity model was tested using non-parametric permutation-based inference (FSL’s randomise tool with 5000 permutations; Winkler *et al*., 2014) with cluster threshold at *Z* = 3.1 (*p* = 0.001) and a *FWE-corrected p* at 0.05.

## RESULTS

Whole-brain multiple regression analysis on the LC connectivity revealed significant regions that have quadratic relationships with age. As expected, a significant positive quadratic relationship of age was found for connectivity between LC and several regions that are mainly associated with the sensory process (i.e., visual, somatosensory, auditory; Figure 1C-E). For example, visual processing regions along the ventral occipitotemporal and dorsal visual pathways including the occipital and temporal fusiform gyrus, parahippocampal gyrus, and precuneus (e.g., Wang *et al*., 2011), decrease functional connectivity with the LC gradually from early childhood years to a low around 40 -45 years old, and then increase into old age (Figure 1C). The parietal operculum extended to the central region and the cerebellum also showed the same u-shape curve of age effects on the LC connectivity (Figure 1D). These regions are known as the secondary somatosensory cortex involved in tactile and pain sensations (Burton *et al*., 2008). Finally, we found that regions in the primary auditory network including *Heschl’*s gyrus extended to the planum temporale (e.g., Warrier *et al*., 2009) showed the same quadratic age relationship on the LC connectivity (Figure 1E). To sum, these results indicated that during their respective development stages children and older adults have increased sensory interaction with the LC in the brain.

Importantly, we also found that there was a significant age-related negative quadratic effect on the LC connectivity for the frontal regions (Figure 1F). The frontal pole extended to the frontal medial cortex, known to be involved in action monitoring and cognitive control (e.g., action selection; Kovach *et al*., 2012), showed lower LC connectivity during the early childhood and older adulthood years than younger adulthood years (i.e., inverted u-shape curve). In other words, the LC has stronger connectivity with the frontal regions during younger adulthood years compared to both developing children and older adults. All significant regions of LC connectivity associated with ages are displayed in Table1 and Figure S2 in *Supplementary Information*.

**Table 1.**
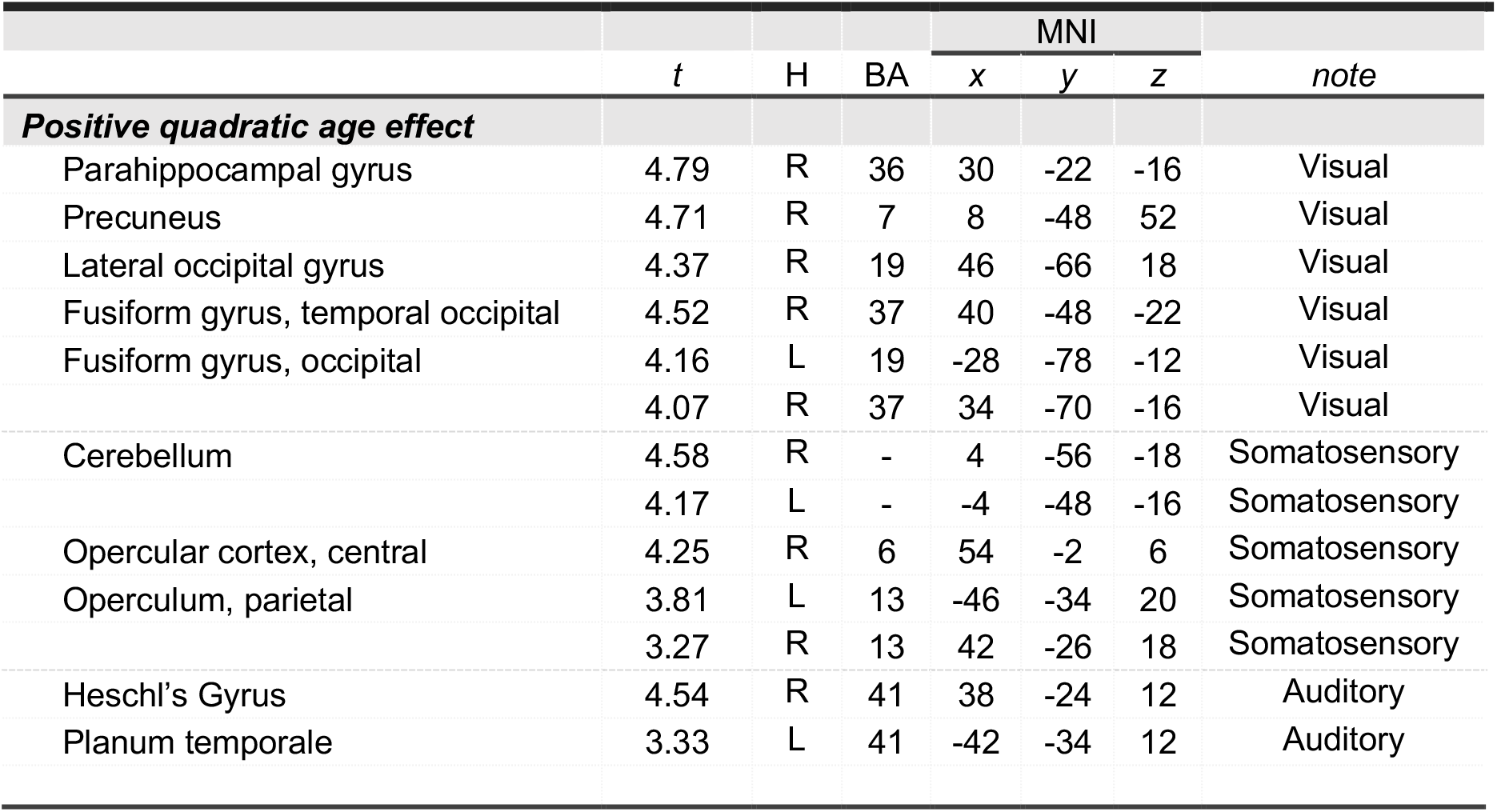
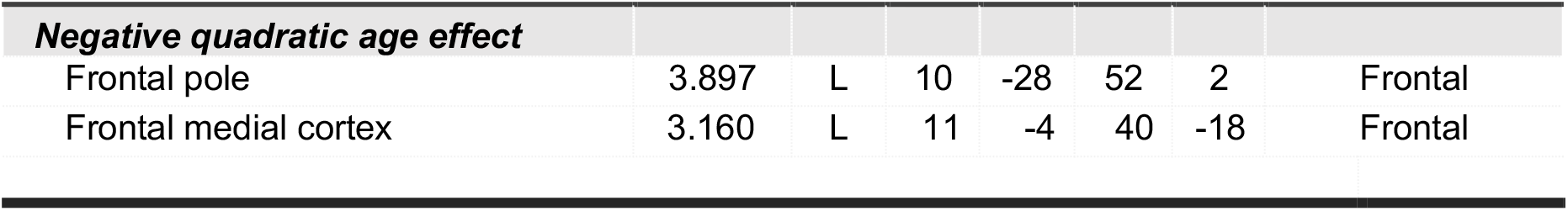
Significant brain regions of quadric age effects (cluster threshold at Z = 3.1 and corrected p value at 0.05 after 5000 permutation) on the LC seed-based whole-brain connectivity analysis. t = t-value; H = hemisphere; BA = Broadman area; Region labeling is based on Harvard-Oxford atlas.

## DISCUSSION

The goal of the current study was to provide a full description of age-related changes in the intrinsic LC connectivity by adopting cross-sectional fMRI data from early childhood to older adulthood. Specifically, given the findings that early children and older adults prone to distraction (Bachman *et al*., 2021; Clewett *et al*., 2016; Kempadoo *et al*., 2016; Liu *et al*., 2020), we hypothesized that the LC, a critical region for selective information processing in the brain, showed the distinct connectivity patterns with other regions in early childhood and older adulthood compared to younger adults who show more stable attentional ability. As a result, we found that the LC’s connectivity with sensory regions showed a u-shaped curve pattern across ages, indicating that the sensory regions exhibit highly increased intrinsic connectivity with the LC in both the early childhood and late older adulthood. Given the previous evidence showing the increased attentional distractibility in both early children and older adults (Bachman *et al*., 2021; Clewett *et al*., 2016; Hoyer *et al*., 2021; Kannass *et al*., 2006; Kempadoo *et al*., 2016; Lee *et al*., 2018; Lee *et al*., 2020c; Leslie *et al*., 1985; Liu *et al*., 2020; Marcyniuk *et al*., 1986; Mueller *et al*., 2017; Olpe and Steinmann, 1982; Palfrey *et al*., 1985), the current findings suggest that such age-related distractibility is possibly due to the insufficient sensory gating process by the LC. Most importantly, the current analyses also revealed that the LC connectivity with the frontal regions showed an inverted u-shape curve pattern. That is, while the sensory network regions are connected to the LC excessively, the frontal network regions have decreased connectivity with the LC, implying that the frontal control regions cannot handle the sensory overflows appropriately. This is the first full description of how the LC configuration changes across ages, informing the LC model of distractibility in both children and older adults.

The current findings implied that the increased distractibility at both early and late developmental stages is due to not only the LC-related excessive sensory overflows but also the lower LC connectivity with the frontal regions. However, although the observed patterns of LC connectivity for sensory and frontal regions are the same for both children and older adult groups, the underlying neural mechanisms for the attentional deficits regarding the LC connectivity may not be the same given structural differences in the developmental trajectory. At the early developmental stage, the primary brain structures including sensory cortex and subcortical bottom-up network regions mature first while the higher-cognitive prefrontal regions are still in the process of developing (Casey *et al*., 2019; Sowell *et al*., 2004), whereas the LC structure is not yet fully developed (W Bondareff *et al*., 1981; William Bondareff *et al*., 1982; Jacobs *et al*., 2018; Liu *et al*., 2019). Thus, it is possible that the LC fails to appropriately prioritize sensory inputs inflow from the fully developed sensory networks, leading to sensory overflows in the children’s brain. That is, the immature LC which cannot control sensory inputs appropriately, and the overflow leads to the unnecessarily increased functional connectivity between LC and sensory regions. In addition, the LC fails to initiate the frontal control region for the flux of sensory inputs in the children’s brain, leading to the decreased functional connectivity. In contrast, given the finding of prominent neurodegenerations of the prefrontal areas compared to other regions (Mattson and Arumugam, 2018), the increased distractibility in elderly is more derived from the decreased frontal functionality in controlling sensory inputs. Although the sensory networks are also functionally degenerated with the frontal cortex in older adults (Salat *et al*., 2004), evidence indicates the decreased sensory sensitivity in the brain at the early sensory gating stage. For instance, the older adults have less activations in the visual and auditory cortex under the passive stimuli presentation (Cliff *et al*., 2013) suggesting that older adults have less sensory-perceptual sensitivity in terms of change detection. Therefore, the LC can still handle the reduced primary sensory processing even when the LC is degenerated, but the prominently decreased frontal control regions are overwhelmed by even less sensory inputs. It possibly represented as the increased functional connectivity between the LC and the sensory and the decreased connectivity between the LC and the frontal control region.

In the current study, we mainly examined the LC-centered neural connectivity across age that possibly serves as the underlying neural mechanism for the attentional distractibility often observed at both early and late developmental stages. However, the suggested LC circuit mechanism sheds lights on understanding other late cognitive process and attention-related mental disorders, as the current result showed the intrinsic connectivity pattern of the LC as a function of age, and it can be used as a framework to interpret the LC-involved neural activities. For instance, the attentional process is involved in memory encoding. Some studies revealed that the LC is associated with memory encoding (Clewett *et al*., 2018; Yebra *et al*., 2019) and older adults with reduced LC structure showed poorer memory encoding (Hämmerer *et al*., 2018). With regard to these studies, our results imply that the intrinsic LC-parahippocampal gyrus connection is a pivotal neural circuit of memory encoding in aging. As an another example, ADHD is regarded as a mental illness characterized by hyperarousal and attentional vigilance (Bruno *et al*., 2007). As described above, it is known that the LC is associated with ADHD. Our results could be helpful to understand and/or predict neural underpinnings of ADHD developmental trajectories given that there have not been many studies involving adults with ADHD (Sudre *et al*., 2017).

However, there are some limitations in the current study. We examined intrinsic functional connectivity of the LC using the non-task based intrinsic neural network (i.e., resting-state fMRI) based on the previous behavioral observations of the increased attentional distractibility in the early childhood and older adulthood. Thus, our observation might be suboptimal to link actual attentional ability and LC associated neural configurations compared to task-based assessments in the laboratory with various attentional tasks, which measure attentional selectivity and control more directly. Future research is needed to employ task-based assessments to link the attentional ability and LC connectivity changes across ages.

Moreover, it is important to note that the LC is an exceptionally small structure in the brainstem, and thus it is difficult to locate its location and signal in an individual brain. Although we used the standard LC structure mask and extracted LC time-series (i.e., LC’s neural activity) from non-smoothed EPI image on the native space (e.g., Alakörkkö *et al*., 2017) combined with the ICA-denoising (Clewett *et al*., 2018; Lee *et al*., 2018; Lee *et al*., 2014) to increase LC signal fidelity in the connectivity estimation, there are several ways to increase LC signal reliability. First, running an additional T1-FSE scan (i.e., neuromelanin sequence-based structural scan) can be used. With a 2-3 minute duration in a scan, it allows to locate the individual specific LC structure on the native space (e.g., Clewett *et al*., 2016). Unfortunately, the current study is based on the public data and thus we could not to utilize additional LC structure images given the pre-determined imaging protocols and collections. Secondly, extracting a seed signal without smoothing on the native space that can minimize the mixture of signals between nearby regions (Lee *et al*., 2020a). In our analysis, we used the non-smoothed LC time-series as a seed region neural activity. Lastly, the LC is often confounded by physio artifacts such as cardiac pulsation and thus it is helpful to run additional physiological noise denoising (Glover *et al*., 2000). Although the ICA-denoising is a promising approach to mitigate physiological influence at the global level, the individual-based physiological denoising process using respiration and cardiac pulse signal can be more focal and direct to the brainstem signal fluctuation correction (Glover *et al*., 2000). In this instance we could not use individual specific LC masks or physiological noise correction. Therefore, in future work, it would be beneficial to utilize the neuromelanin sequence LC structural scan and physiological data collection.

## Author Contribution

Conceptualization, I.S. and T.-H.L.; methodology, T.-H.L.; software, I.S.; validation, I.S., J.N. and T.-H.L.; formal analysis, T.-H.L.; investigation, I.S. and T.-H.L.; resources, J.N. and T.-H.L.; data curation, I.S.; writing—original draft preparation, I.S. and T.-H.L.; writing—review and editing, J.N. and T.-H.L.; visualization, I.S.; supervision, T.-H.L.; project administration, T.-H.L.; funding acquisition, T.-H.L. All authors have read and agreed to the published version of the manuscript.

## Institutional Review Board Statement

All data were acquired in accordance with Nathan-Kline-Institute IRB guidelines (Phase I #226781 and Phase II #239708).

## REFERENCES

Alakörkkö, T., Saarimäki, H., Glerean, E., Saramäki, J., and Korhonen, O. (2017). Effects of spatial smoothing on functional brain networks. European Journal of Neuroscience 46, 2471–2480.

Aston-Jones, G., and Cohen, J.D. (2005). An integrative theory of locus coeruleus-norepinephrine function: adaptive gain and optimal performance. Annu. Rev. Neurosci. 28, 403–450.

Avants, B.B., Tustison, N.J., Stauffer, M., Song, G., Wu, B., and Gee, J.C. (2014). The Insight ToolKit image registration framework. Frontiers in neuroinformatics 8, 44.

Bachman, S.L., Dahl, M.J., Werkle-Bergner, M., Düzel, S., Forlim, C.G., Lindenberger, U., Kühn, S., and Mather, M. (2021). Locus coeruleus MRI contrast is associated with cortical thickness in older adults. Neurobiology of Aging 100, 72–82.

Berridge, C.W., and Waterhouse, B.D. (2003). The locus coeruleus-noradrenergic system: modulation of behavioral state and state-dependent cognitive processes. Brain Res. Brain Res. Rev. 42, 33–84. 10.1016/s0165-0173(03)00143-7.

Bland, A.R. (2012). Different varieties of uncertainty in human decision-making. Frontiers in neuroscience 6, 85.

Bondareff, W., Mountjoy, C., and Roth, M. (1981). Selective loss of neurones of origin of adrenergic projection to cerebral cortex (nucleus locus coeruleus) in senile dementia. The Lancet 317, 783–784.

Bondareff, W., Mountjoy, C.Q., and Roth, M. (1982). Loss of neurons of origin of the adrenergic projection to cerebral cortex (nucleus locus ceruleus) in senile dementia. Neurology 32, 164–164.

Bruno, K.J., Freet, C.S., Twining, R.C., Egami, K., Grigson, P.S., and Hess, E.J. (2007). Abnormal latent inhibition and impulsivity in coloboma mice, a model of ADHD. Neurobiology of disease 25, 206–216.

Burton, H., Sinclair, R.J., Wingert, J.R., and Dierker, D.L. (2008). Multiple parietal operculum subdivisions in humans: tactile activation maps. Somatosensory & motor research 25, 149–162.

Carter, M.E., Yizhar, O., Chikahisa, S., Nguyen, H., Adamantidis, A., Nishino, S., Deisseroth, K., and de Lecea, L. (2010). Tuning arousal with optogenetic modulation of locus coeruleus neurons. Nat Neurosci 13, 1526-1533. 10.1038/nn.2682.

Casey, B., Heller, A.S., Gee, D.G., and Cohen, A.O. (2019). Development of the emotional brain. Neuroscience letters 693, 29–34.

Chen, W. (2002). Stimulant drugs and ADHD: Basic and clinical neuroscience. edited by M. Solanto, A. Arnsten, & F.X. Castellanos. University of Oxford Press, Oxford, 2001. pp. 410. US$79.95 (hb). ISBN: 0-19-513371-4. Journal of Child Psychology and Psychiatry 43, 413–414. 10.1111/1469-7610.0032b.

Chiew, K.S., and Braver, T.S. (2013). Temporal dynamics of motivation-cognitive control interactions revealed by high-resolution pupillometry. Frontiers in psychology 4, 15.

Clewett, D.V., Huang, R., Velasco, R., Lee, T.-H., and Mather, M. (2018). Locus Coeruleus Activity Strengthens Prioritized Memories Under Arousal. J Neurosci 38, 1558-1574. 10.1523/JNEUROSCI.2097-17.2017.

Clewett, D.V., Lee, T.-H., Greening, S., Ponzio, A., Margalit, E., and Mather, M. (2016). Neuromelanin marks the spot: identifying a locus coeruleus biomarker of cognitive reserve in healthy aging. Neurobiol. Aging 37, 117–126. 10.1016/j.neurobiolaging.2015.09.019.

Cliff, M., Joyce, D.W., Lamar, M., Dannhauser, T., Tracy, D.K., and Shergill, S.S. (2013). Aging effects on functional auditory and visual processing using fMRI with variable sensory loading. Cortex 49, 1304–1313.

Coronel-Oliveros, C., Castro, S., Cofré, R., and Orio, P. (2021). Structural features of the human connectome that facilitate the switching of brain dynamics via noradrenergic neuromodulation. Frontiers in Computational Neuroscience 15, 61.

Darcq, E., and Kieffer, B.L. (2015). PI 3K signaling in the locus coeruleus: a new molecular pathway for ADHD research. EMBO molecular medicine 7, 859–861.

Devilbiss, D.M., and Waterhouse, B.D. (2011). Phasic and tonic patterns of locus coeruleus output differentially modulate sensory network function in the awake rat. Journal of neurophysiology 105, 69–87.

Devilbiss, D.M., Waterhouse, B.D., Berridge, C.W., and Valentino, R. (2012). Corticotropin-releasing factor acting at the locus coeruleus disrupts thalamic and cortical sensory-evoked responses. Neuropsychopharmacology 37, 2020–2030.

Elliott, M.L., Knodt, A.R., Cooke, M., Kim, M.J., Melzer, T.R., Keenan, R., Ireland, D., Ramrakha, S., Poulton, R., and Caspi, A. (2019). General functional connectivity: Shared features of resting-state and task fMRI drive reliable and heritable individual differences in functional brain networks. Neuroimage 189, 516–532.

Fast, C.D., and McGann, J.P. (2017). Amygdalar gating of early sensory processing through interactions with locus coeruleus. Journal of Neuroscience 37, 3085–3101.

Gallant, S.N., Kennedy, B.L., Bachman, S.L., Huang, R., Lee, T.-H., and Mather, M. (2021). Behavioral and fMRI evidence that arousal enhances bottom-up attention and memory selectivity in young but not older adults. bioRxiv.

Glover, G.H., Li, T.Q., and Ress, D. (2000). Image-based method for retrospective correction of physiological motion effects in fMRI: RETROICOR. Magn. Reson. Med. 44, 162–167. 10.1002/1522-2594(200007)44:1<162::aid-mrm23>3.0.co;2-e.

Hoyer, R.S., Elshafei, H., Hemmerlin, J., Bouet, R., and Bidet-Caulet, A. (2021). Why are children so distractible? Development of attention and motor control from childhood to adulthood. Child development.

Hämmerer, D., Callaghan, M.F., Hopkins, A., Kosciessa, J., Betts, M., Cardenas-Blanco, A., Kanowski, M., Weiskopf, N., Dayan, P., Dolan, R.J., and Düzel, E. (2018). Locus coeruleus integrity in old age is selectively related to memories linked with salient negative events. Proceedings of the National Academy of Sciences 115, 2228–2233. 10.1073/pnas.1712268115.

Jacobs, H.I., Müller-Ehrenberg, L., Priovoulos, N., and Roebroeck, A. (2018). Curvilinear locus coeruleus functional connectivity trajectories over the adult lifespan: a 7T MRI study. Neurobiology of aging 69, 167–176.

Kannass, K.N., Oakes, L.M., and Shaddy, D.J. (2006). A longitudinal investigation of the development of attention and distractibility. Journal of Cognition and Development 7, 381–409.

Kempadoo, K.A., Mosharov, E.V., Choi, S.J., Sulzer, D., and Kandel, E.R. (2016). Dopamine release from the locus coeruleus to the dorsal hippocampus promotes spatial learning and memory. Proc. Natl. Acad. Sci. U. S. A. 113, 14835-14840. 10.1073/pnas.1616515114.

Keren, N.I., Lozar, C.T., Harris, K.C., Morgan, P.S., and Eckert, M.A. (2009). In vivo mapping of the human locus coeruleus. Neuroimage 47, 1261–1267.

Kovach, C.K., Daw, N.D., Rudrauf, D., Tranel, D., O’Doherty, J.P., and Adolphs, R. (2012). Anterior prefrontal cortex contributes to action selection through tracking of recent reward trends. Journal of Neuroscience 32, 8434–8442.

Landis, D., Courtney, W., Dieringer, C., Kelly, R., King, M., Miller, B., Wang, R., Wood, D., Turner, J.A., and Calhoun, V.D. (2016). COINS Data Exchange: An open platform for compiling, curating, and disseminating neuroimaging data. NeuroImage 124, 1084–1088.

Lee, T.-H., Kim, S.H., Katz, B., and Mather, M. (2020a). The Decline in Intrinsic Connectivity Between the Salience Network and Locus Coeruleus in Older Adults: Implications for Distractibility. Frontiers in Aging Neuroscience 12. 10.3389/fnagi.2020.00002.

Lee, T.-H., Miernicki, M.E., and Telzer, E.H. (2017a). Behavioral and neural concordance in parent-child dyadic sleep patterns. Dev. Cogn. Neurosci. 26, 77–83. 10.1016/j.dcn.2017.06.003.

Lee, T.-H., Miernicki, M.E., and Telzer, E.H. (2017b). Families that fire together smile together: Resting state connectome similarity and daily emotional synchrony in parent-child dyads. Neuroimage 152, 31-37. 10.1016/j.neuroimage.2017.02.078.

Lee, T.-H., and Telzer, E.H. (2016). Negative functional coupling between the right fronto-parietal and limbic resting state networks predicts increased self-control and later substance use onset in adolescence. Dev. Cogn. Neurosci. 20, 35-42. 10.1016/j.dcn.2016.06.002.

Lee, T.-H., Greening, S.G., Ueno, T., Clewett, D. V., Ponzio, A., Sakaki, M., and Mather, M. (2018). Arousal increases neural gain via the locus coeruleus-norepinephrine system in younger adults but not in older adults. Nat Hum Behav 2, 356-366. 10.1038/s41562-018-0344-1.

Lee, T.-H., Kim, S.H., Katz, B., and Mather, M. (2020b). The Decline in Intrinsic Connectivity Between the Salience Network and Locus Coeruleus in Older Adults: Implications for Distractibility. Front Aging Neurosci 12, 2. 10.3389/fnagi.2020.00002.

Lee, T.-H., Kim, S.H., Katz, B., and Mather, M. (2020c). The Decline in Intrinsic Connectivity Between the Salience Network and Locus Coeruleus in Older Adults: Implications for Distractibility. Frontiers in Aging Neuroscience 12. 10.3389/fnagi.2020.00002.

Lee, T.-H., Sakaki, M., Cheng, R., Velasco, R., and Mather, M. (2014). Emotional arousal amplifies the effects of biased competition in the brain. Soc Cogn Affect Neurosci 9, 2067–2077. 10.1093/scan/nsu015.

Leslie, F.M., Loughlin, S.E., Sternberg, D.B., McGaugh, J.L., Young, L.E., and Zornetzer, S.F. (1985). Noradrenergic changes and memory loss in aged mice. Brain research 359, 292–299.

Liu, K.Y., Acosta-Cabronero, J., Cardenas-Blanco, A., Loane, C., Berry, A.J., Betts, M.J., Kievit, R.A., Henson, R.N., Düzel, E., and Howard, R. (2019). In vivo visualization of age-related differences in the locus coeruleus. Neurobiology of aging 74, 101–111.

Liu, K.Y., Kievit, R.A., Tsvetanov, K.A., Betts, M.J., Düzel, E., Rowe, J.B., Howard, R., and Hämmerer, D. (2020). Noradrenergic-dependent functions are associated with age-related locus coeruleus signal intensity differences. Nature communications 11, 1–9.

Marcyniuk, B., Mann, D., and Yates, P. (1986). Loss of nerve cells from locus coeruleus in Alzheimer’s disease is topographically arranged. Neuroscience letters 64, 247–252.

Mather, M., and Harley, C.W. (2016). The Locus Coeruleus: Essential for Maintaining Cognitive Function and the Aging Brain. Trends Cogn Sci 20, 214–226. 10.1016/j.tics.2016.01.001.

Mattson, M.P., and Arumugam, T.V. (2018). Hallmarks of Brain Aging: Adaptive and Pathological Modification by Metabolic States. Cell Metab 27, 1176–1199. 10.1016/j.cmet.2018.05.011.

McBurney-Lin, J., Lu, J., Zuo, Y., and Yang, H. (2019). Locus coeruleus-norepinephrine modulation of sensory processing and perception: A focused review. Neuroscience & Biobehavioral Reviews 105, 190–199.

McCall, J.G., Siuda, E.R., Bhatti, D.L., Lawson, L.A., McElligott, Z.A., Stuber, G.D., and Bruchas, M.R. (2017). Locus coeruleus to basolateral amygdala noradrenergic projections promote anxiety-like behavior. Elife 6, e18247.

Mueller, A., Hong, D.S., Shepard, S., and Moore, T. (2017). Linking ADHD to the neural circuitry of attention. Trends in cognitive sciences 21, 474–488.

Naegeli, C., Zeffiro, T., Piccirelli, M., Jaillard, A., Weilenmann, A., Hassanpour, K., Schick, M., Rufer, M., Orr, S.P., and Mueller-Pfeiffer, C. (2018). Locus coeruleus activity mediates hyperresponsiveness in posttraumatic stress disorder. Biological psychiatry 83, 254–262.

Nomi, J.S., Bolt, T.S., Ezie, C.C., Uddin, L.Q., and Heller, A.S. (2017). Moment-to-moment BOLD signal variability reflects regional changes in neural flexibility across the lifespan. Journal of Neuroscience 37, 5539–5548.

Nooner, K.B., Colcombe, S., Tobe, R., Mennes, M., Benedict, M., Moreno, A., Panek, L., Brown, S., Zavitz, S., and Li, Q. (2012). The NKI-Rockland sample: a model for accelerating the pace of discovery science in psychiatry. Frontiers in neuroscience 6, 152.

Olpe, H.-R., and Steinmann, M.W. (1982). Age-related decline in the activity of noradrenergic neurons of the rat locus coeruleus. Brain Research 251, 174–176.

Palfrey, J.S., Levine, M.D., Walker, D.K., and Sullivan, M. (1985). The emergence of attention deficits in early childhood: a prospective study. Journal of Developmental and Behavioral Pediatrics.

Pruim, R.H., Mennes, M., van Rooij, D., Llera, A., Buitelaar, J.K., and Beckmann, C.F. (2015). ICA-AROMA: A robust ICA-based strategy for removing motion artifacts from fMRI data. Neuroimage 112, 267–277.

Rodenkirch, C., Liu, Y., Schriver, B.J., and Wang, Q. (2019). Locus coeruleus activation enhances thalamic feature selectivity via norepinephrine regulation of intrathalamic circuit dynamics. Nature neuroscience 22, 120–133.

Rowe, D., Robinson, P., and Gordon, E. (2005). Stimulant drug action in attention deficit hyperactivity disorder (ADHD): inference of neurophysiological mechanisms via quantitative modelling. Clinical neurophysiology 116, 324–335.

Salat, D.H., Buckner, R.L., Snyder, A.Z., Greve, D.N., Desikan, R.S., Busa, E., Morris, J.C., Dale, A.M., and Fischl, B. (2004). Thinning of the cerebral cortex in aging. Cereb Cortex 14, 721–730. 10.1093/cercor/bhh032.

Serova, L.I., Nwokafor, C., Van Bockstaele, E.J., Reyes, B.A., Lin, X., and Sabban, E.L. (2019). Single prolonged stress PTSD model triggers progressive severity of anxiety, altered gene expression in locus coeruleus and hypothalamus and effected sensitivity to NPY. European Neuropsychopharmacology 29, 482–492.

Sowell, E.R., Thompson, P.M., and Toga, A.W. (2004). Mapping changes in the human cortex throughout the span of life. Neuroscientist 10, 372-392. 10.1177/1073858404263960.

Sudre, G., Szekely, E., Sharp, W., Kasparek, S., and Shaw, P. (2017). Multimodal mapping of the brain’s functional connectivity and the adult outcome of attention deficit hyperactivity disorder. Proc Natl Acad Sci U S A 114, 11787-11792. 10.1073/pnas.1705229114.

Toga, A.W., Thompson, P.M., and Sowell, E.R. (2006). Mapping brain maturation. Focus 29, 148–390.

van der Wel, P., and van Steenbergen, H. (2018). Pupil dilation as an index of effort in cognitive control tasks: A review. Psychonomic bulletin & review 25, 2005–2015.

Wang, Q., Gao, E., and Burkhalter, A. (2011). Gateways of ventral and dorsal streams in mouse visual cortex. Journal of Neuroscience 31, 1905–1918.

Warrier, C., Wong, P., Penhune, V., Zatorre, R., Parrish, T., Abrams, D., and Kraus, N. (2009). Relating structure to function: Heschl’s gyrus and acoustic processing. Journal of Neuroscience 29, 61–69.

Waterhouse, B.D., and Navarra, R.L. (2019). The locus coeruleus-norepinephrine system and sensory signal processing: A historical review and current perspectives. Brain research 1709, 1–15.

Winkler, A.M., Ridgway, G.R., Webster, M.A., Smith, S.M., and Nichols, T.E. (2014). Permutation inference for the general linear model. Neuroimage 92, 381–397.

Yebra, M., Galarza-Vallejo, A., Soto-Leon, V., Gonzalez-Rosa, J.J., de Berker, A.O., Bestmann, S., Oliviero, A., Kroes, M.C.W., and Strange, B.A. (2019). Action boosts episodic memory encoding in humans via engagement of a noradrenergic system. Nat Commun 10, 3534. 10.1038/s41467-019-11358-8.

Zerbi, V., Floriou-Servou, A., Markicevic, M., Vermeiren, Y., Sturman, O., Privitera, M., von Ziegler, L., Ferrari, K.D., Weber, B., De Deyn, P.P., et al. (2019). Rapid Reconfiguration of the Functional Connectome after Chemogenetic Locus Coeruleus Activation. Neuron 103, 702-718.e705. 10.1016/j.neuron.2019.05.034.

