## Supplemental Figure1 & 2 for "Age-related intrinsic functional connectivity changes of locus coeruleus from childhood to older adults"

**SUPPLEMENTARY MATERIALS**

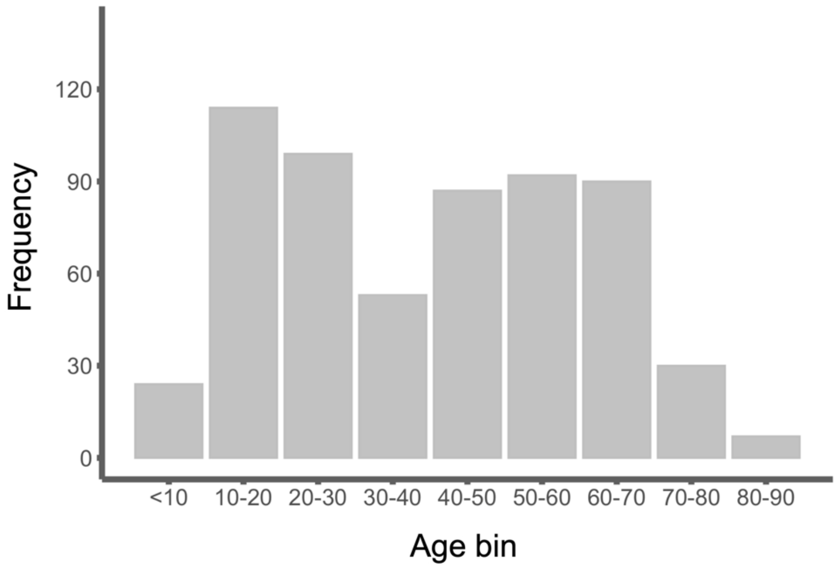


**Figure S1.** Age distribution

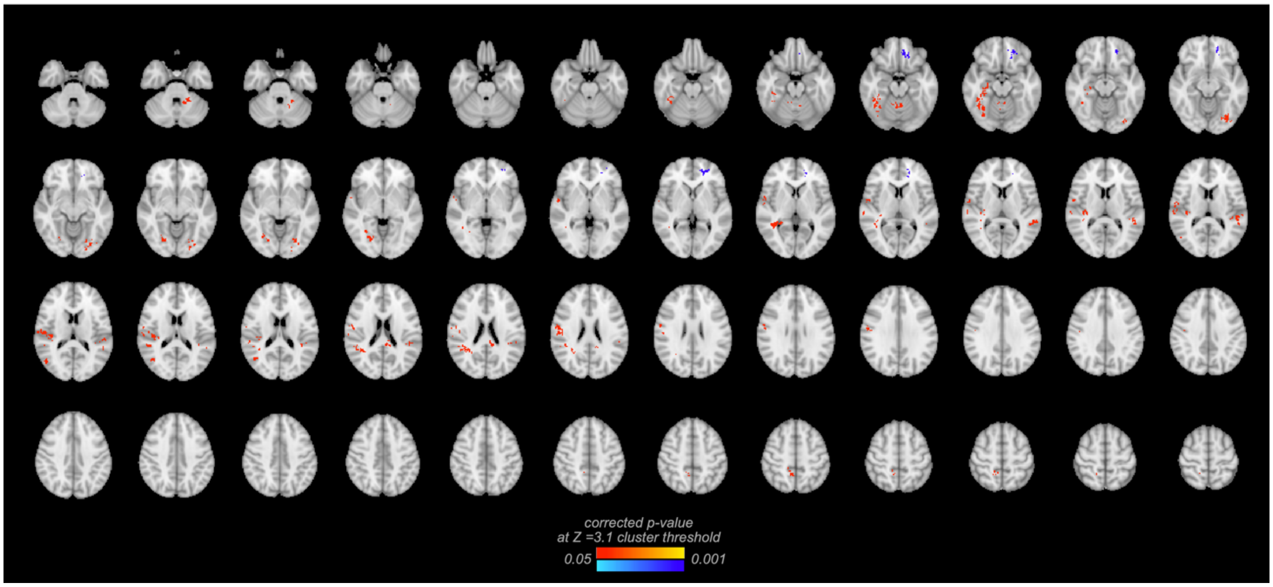


**Figure S2.** The whole-brain map of LC seed-based functional connectivity showing the quadratic effects of age.
